# Trace-based correction of breathing-induced field fluctuations in T2^*^-weighted imaging of the spinal cord

**DOI:** 10.1101/406371

**Authors:** S. Johanna Vannesjo, Stuart Clare, Lars Kasper, Irene Tracey, Karla L. Miller

**Affiliations:** Wellcome Centre for Integrative Neuroimaging, FMRIB, Nuffield Department of Clinical Neurosciences, University of Oxford, Oxford, UK; Institute for Biomedical Engineering, ETH Zurich and University of Zurich, Zurich, Switzerland; Translational Neuromodeling Unit, Institute for Biomedical Engineering, University of Zurich and ETH Zurich, Zurich, Switzerland

**Keywords:** Spinal cord imaging, 7T MRI, Breathing-induced field fluctuations, T2* mapping, Multi-shot EPI

## Abstract

**Purpose:** Spinal cord MRI at ultra-high field is hampered by time-varying magnetic fields associated with the breathing cycle, giving rise to ghosting artifacts in multi-shot acquisitions. Here, we suggest a correction approach based on linking the signal from a respiratory bellows to field changes inside the spinal cord. The information is used to correct the data at the image reconstruction level.

**Methods:** The correction was demonstrated in the context of multi-shot T2*-weighted imaging of the cervical spinal cord at 7T. A respiratory trace was acquired during a high-resolution multi-echo gradient-echo sequence, used for structural imaging and quantitative T2* mapping, and a multi-shot EPI time series, as would be suitable for fMRI. The coupling between the trace and the breathing-induced fields was determined by a short calibration scan in each individual. Images were reconstructed with and without trace-based correction.

**Results:** In the multi-echo GRE, breathing-induced fields caused severe ghosting in the long echo time images, which led to a systematic underestimation of T2* in the spinal cord. The trace-based correction reduced the ghosting and increased the estimated T2* values. Breathing-related ghosting was also observed in the multi-shot EPI images. The correction largely removed the ghosting, thereby improving the temporal signal-to-noise ratio of the time series.

**Conclusions:** Trace-based retrospective correction of breathing-induced field variations can reduce ghosting and improve quantitative metrics in multi-shot structural and functional T2*-weighted imaging of the spinal cord. The method is straightforward to implement and does not rely on sequence modifications or additional hardware beyond a respiratory bellows.

## Introduction

Spatial encoding in MRI relies on the assumption that the background magnetic field is homogenous and stable over time. However, the presence of a subject in the scanner gives rise to both static field inhomogeneity and dynamic field fluctuations. Magnetic susceptibility differences between tissue and air, cause local field distortions around interfaces (1). Because breathing changes the air-tissue distribution of the thorax and abdomen, the surrounding field distribution varies periodically with the breathing cycle (2). The time-varying fields cause mis-localisation of signal, which can manifest as image distortion, apparent motion, blurring or ghosting, depending on the sequence. In neuroimaging, breathing-related field fluctuations can be measured as far away as in the brain (3), but are particularly prominent in the spine because of the proximity to the lungs (4,5). Indeed, breathing-induced fields have been identified as one of the major challenges to overcome to achieve robust functional MRI (fMRI) of the spinal cord (6).

One way to address the problem is to track the field variations over time, and use the information for prospective or retrospective correction approaches. One potential tracker is the signal from a respiratory bellows, which indicates the state of the breathing cycle. It has recently been shown that the respiratory trace can predict over 90% of the breathing-induced time-variance of the field in the spinal cord during normal shallow breathing (5). A respiratory trace has previously been used as basis for real-time shim updating in ultra-high field brain imaging (7), and has more recently been explored for real-time shimming of the spinal cord at 3T using a custom-built 24-channel shim coil (8,9). However, real-time shim updating demands specialized hardware, which is not available at most sites.

In this work, we investigate using the signal from a respiratory bellows to retrospectively correct the acquired MR data for breathing-induced field fluctuations in spinal cord imaging. A linear model is used to transform the acquired respiratory trace to field variations inside the spine along the superior-inferior (z) axis. The model parameters are determined in each individual subject using a short calibration set of fast phase-sensitive FLASH acquisitions. The breathing-induced field variations are then estimated from the trace alone during subsequent scans, and the acquired MR signal is demodulated by the corresponding phase variations before image reconstruction. The method can be implemented on standard MR systems, and for any type of sequence. Here we explore the correction in the context of T2*-weighted imaging at 7T. Ultra-high field accentuates the T2*-weighted contrast and allows for higher resolution. However, T2*-weighted acquisitions are particularly vulnerable to the effects of field fluctuations. Furthermore, the amplitude of the field variations scale with the strength of the background field. Hence, accurate correction is crucial for optimal image quality in ultra-high field T2*-weighted imaging. While T2* is therefore challenging at ultra-high field, it is also of particular interest in the spine. In structural imaging, T2*-weighting provides excellent gray/white matter contrast in the spine (10–12) and it is also the contrast underlying blood-oxygen-level-dependent functional imaging. We therefore implement the correction for high-resolution multi-echo gradient-echo acquisitions, used for structural imaging and quantitative T2* mapping, and time series of multi-shot EPI images, intended for functional imaging. We focus on multi-shot acquisitions as they are more robust against static B0 field inhomogeneity compared to single-shot acquisitions, while being especially affected by dynamic field variations.

## Methods

All acquisitions were performed on a whole-body 7T system (Magnetom, Siemens Healthineers, Erlangen, Germany). Imaging was performed with a volume-transmit,16-channel receive cervical spine coil (Quality Electrodynamics, Mayfield village, OH, USA) in eight healthy volunteers (1 female, mean (range) age 34 (27-40) years, weight 79 (55-95) kg, height 1.81 (1.64-1.93) m, body mass index 23.8 (20.4-29.0) kg/m^2^), in compliance with local ethics guidelines. The volunteers were instructed to breathe regularly at a comfortable pace and to avoid swallowing during the scans. During all acquisitions, the signal from a respiratory bellows placed just below the thorax was acquired. The respiratory trace, *R(t)*, was synchronized with the imaging data via a trigger in the sequence. No low-pass filter was applied to the trace, but the mean offset was subtracted for each scan to remove slow baseline drifts between scans.

### Trace-based correction

The trace-based correction pipeline is summarized in Figure 1. Calibration of the individual breathing-induced spatial field profiles was performed as described in Vannesjo et al (5). In brief, a time series of FLASH images (13) (resolution 3.4×2.3×3.0 mm^3^, FOV 144×144 mm^2^, TR 8 ms, TE 4.08 ms, bandwidth 240 Hz/pixel, FA 6 ˚) of a single sagittal slice through the center of the spinal cord was acquired during normal breathing (Fig. 1a). The volume TR of the acquisition was 344 ms, yielding sufficient temporal resolution to capture the breathing cycle. The phase in each voxel, *ϕ(r,t)*, was unwrapped over time and the time-averaged phase, 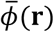, was subtracted. The resulting phase evolution was divided by the echo time to yield a measure of the field changes over time: 

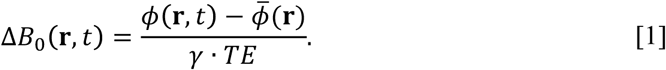

**Figure 1:**
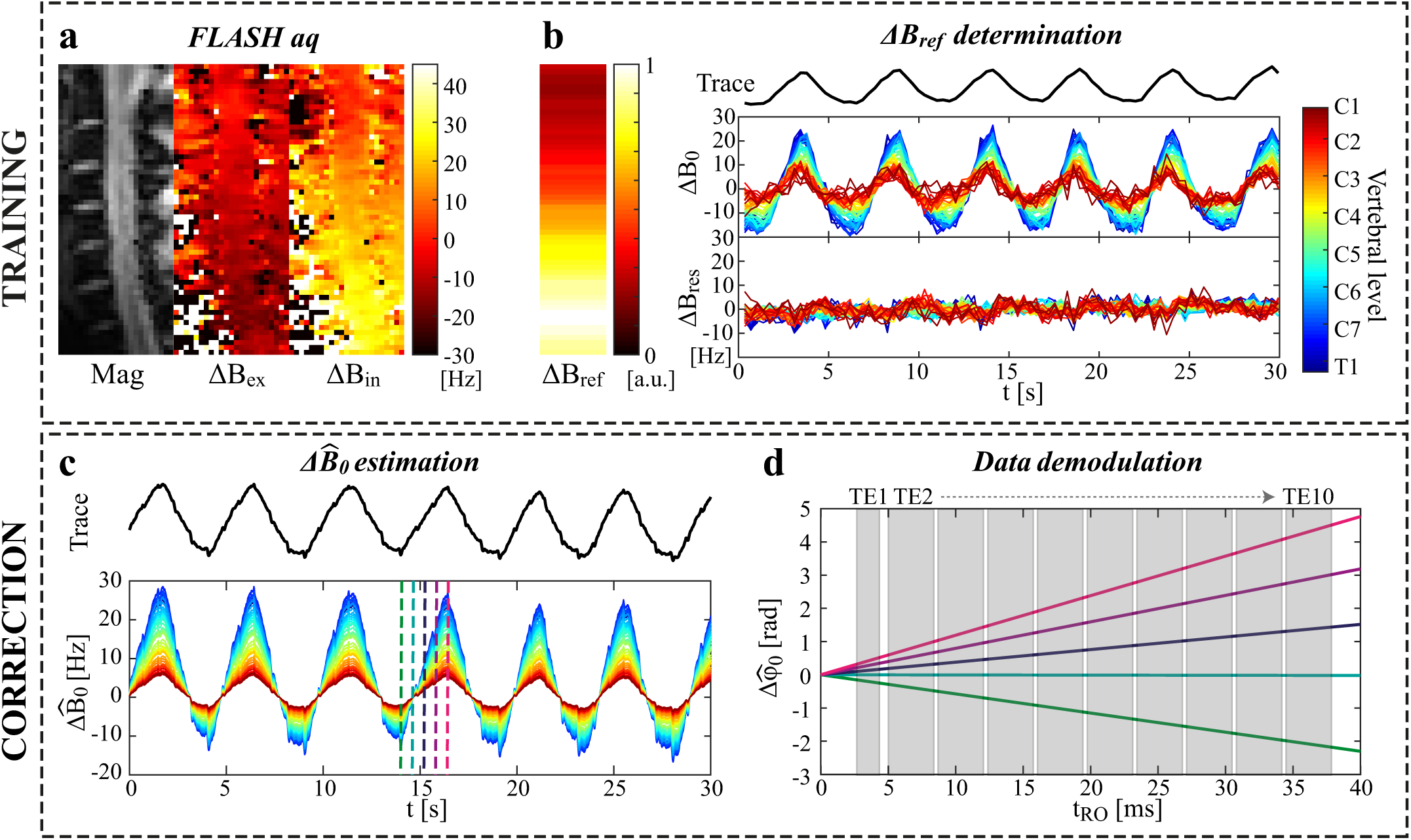
Schematic of the trace-based correction method. **a)** Magnitude and phase images of a single sagittal slice are acquired with a FLASH sequence. The phase images are here shown at a peak of expiration (Δ*B*_*ex*_) and inspiration (ΔB_in_). **b)** The coupling parameter (ΔB_ref_(z)) between the respiratory trace and the field offset (δB_0_(z,t)) inside the spinal cord is determined for each axial slice based on the FLASH calibration data. The residual field offsets (ΔB_ref_(z,t)) show that the linear model explains a large part of the measured temporal field variations. **c)** During later scans, the field offset (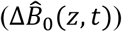) can be estimated based on ΔB_ref_(z)and the respiratory trace. **d)** The estimated field offset yields corresponding phase evolutions (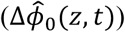), here shown for a multi-echo GRE readout at five different time points indicated by vertical lines in the plot in c. The acquired image data is demodulated by the estimated 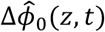.

A mask covering the spinal cord was manually defined on the FLASH magnitude image. The field within the mask was averaged in the transverse plane, yielding a time-series of measured field offsets, δB_0_(z,t), along the superior-inferior (z) axis (Fig. 1b). A linear model linking the respiratory trace *R(t)* to the estimated field offset, 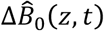, at any given z location was assumed:

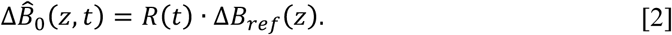

A least-squares fit based on 30 seconds of the calibration data was used to determine Δ*B*_*ref*_ (*z*) from Eq. [2] in each subject. Time points affected by swallowing were excluded from the fit. The measured Δ*B*_*ref*_ (*z*) was then used to estimate the field fluctuations from the respiratory trace acquired during subsequent scans (Fig. 1c).

For the correction, a spatially homogenous field offset within each transverse plane was assumed. The field offset was assumed to be static during the length of the readout train. In this case the MR signal from the object is modulated by a phase offset, Δ*ϕ*_0_, given by (Fig. 1d):

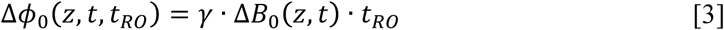

where *t*_*R0*_ denotes time since the last RF excitation, and *t* represents time over the full length of the MR sequence. The acquired imaging data, *s(z,t,t*_*R0*_), was demodulated with the estimated phase offset, 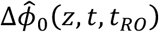, for each time point in the acquisition:

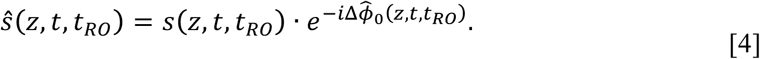

After demodulation, image reconstruction was performed with an iterative conjugate gradient optimization algorithm with SENSE unfolding (14). For comparison, image reconstruction was also performed without prior demodulation of the phase offset, to yield uncorrected images.

### Acquisitions

Data for correction were acquired using two sequences: a multi-echo GRE sequence using a single-line readout at each echo time, suitable for T2*-weighted structural imaging and quantitative T2* mapping, and a single-TE GRE sequence using a segmented EPI readout, suitable for high-resolution functional imaging.

The multi-echo GRE sequence had the following imaging parameters: 24 axial slices, resolution 0.35×0.35×3 mm^3^, FOV 146×128 mm^2^, TR 1000 ms, 10 echoes: TE [3.51, 6.68, 10.37, 14.06, 17.75, 21.44, 25.13, 28.82, 32.51, 36.20] ms, bipolar readout, bandwidth 600 Hz/pixel for the first echo and 300 Hz/pixel for all later echoes, AP phase encoding, flip angle 46 ˚, T_acq_ 6:06 min. The increased bandwidth of the first echo served to minimise the first TE in order to achieve close to proton density contrast. The acquisitions were centered on the lower end of the C4 vertebra to fully cover the C3-C6 vertebral levels. The magnitude images of the separate echoes (with or without correction) were combined with a root-sum-of-squares (RSS) combination to form high-resolution structural images. Quantitative T2*-mapping was performed by a voxel-wise fit of a mono-exponential function to the single-echo images. The fit was performed in Matlab 2017a using the Trust-Region algorithm of the ‘fit’ function. For evaluation of the spinal cord T2* mapping, a mask covering the spinal cord was created on the RSS images with a semi-automatic approach using the ‘sct_propseg’ function of the Spinal Cord Toolbox (15,16). Starting points for identifying the spinal cord centerline were selected manually.

The multi-shot EPI sequence (24 axial slices, resolution 0.76×0.76×3 mm, FOV 128×128 mm, SENSE factor 2, 4 shots, TR 650 ms, volume TR 2.6 s, TE 14 ms, bandwidth 1144 Hz/pixel, AP phase encoding, flip angle 42 ˚, 120 volumes, T_acq_ 5:17 min) was acquired in six of the volunteers (1 female, mean (range) age 34 (28-40) years, weight 78 (55-95) kg, height 1.82 (1.64-1.93) m, body mass index 23.3 (20.4-26.9) kg/m_^2^_). In order to minimize the achievable TE in light of very short T2* in the spine, the EPI sequences did not include phase correction lines for static ghost removal; this correction was performed using corresponding phase correction lines acquired separately in a phantom. A mask covering the spinal cord was extracted for the multi-shot EPI data, as described above. The temporal signal-to-noise ratio (tSNR) was calculated inside the spinal cord mask with and without correction.

## Results

Figure 2 shows single echoes and the RSS echo combination of the high-resolution structural acquisition in one subject. In the uncorrected single echo images, there is an irregular ghosting pattern, which increases with echo time. The ghosting smears out the signal from the spinal cord over the image, to the degree that the depiction of the spinal cord is completely lost in later echoes in some slices. The artefacts are generally more severe towards lower levels of the cervical spine, where the field fluctuations are higher. In the RSS images, the ghosting results in a blurred appearance, reduced signal amplitude, and diminished gray/white matter contrast. With correction, the ghosting is reduced, yielding more sharply delineated structures and improved gray/white matter contrast.

**Figure 2:**
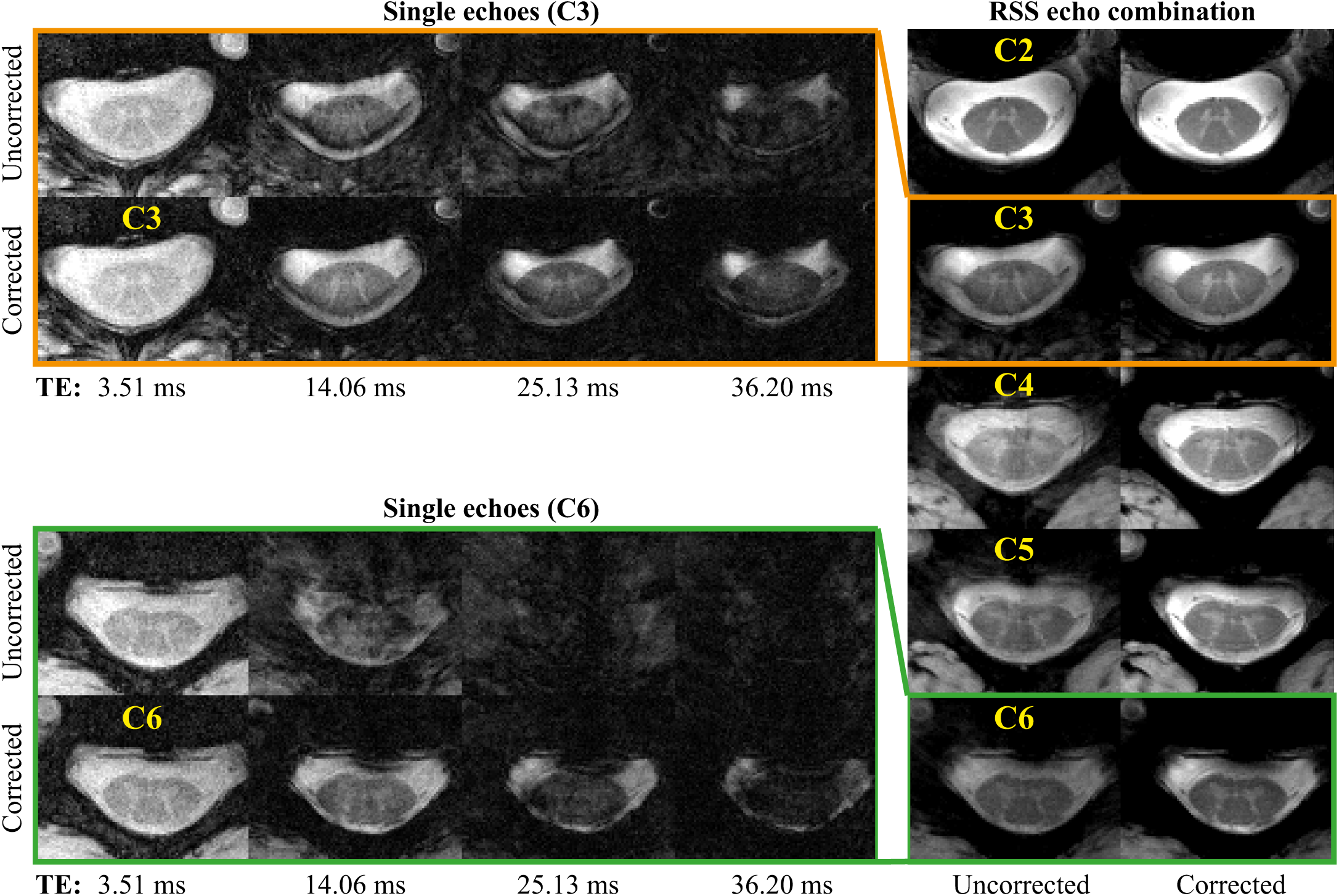
Multi-echo gradient echo acquisition in one subject. To the left are single echo images from the 1^st^, 4^th^, 7^th^ and 10^th^ echo at two different vertebral levels (C3 and C6), without and with correction. To the right are the root-sum-of-squares (RSS) echo combination images shown at vertebral levels C2-C6, without and with correction.

The ghosting systematically affects the T2* quantification as shown in Figure 3. The upper half of the plot (Fig 3a-b) shows the quantification results in two different slices from one subject. The mean signal within the spinal cord decays faster in the uncorrected case (Fig. 3b), as the ghosting increasingly scatters the signal from the spinal cord at longer echo times. This results in a systematic under-estimation of the local T2*, as displayed in the T2*-maps (Fig. 3a) and the histogram of measured T2* values within the spinal cord mask (Fig. 3b). All subjects showed a systematic shift towards higher T2* values inside the spinal cord with the correction (Fig. 3c). The median T2* value within the spinal cord mask was 15-23 ms in the uncorrected case and 23-35 ms with correction (Fig. 3d). In locations where the T2* was intrinsically low, the correction did not affect the quantification. This is evident from the upper slice in Figure 3a, where local static field inhomogeneity due to close proximity to the C2/C3 intervertebral junction caused a marked shortening of T2* along the dorsal edge of the spinal cord (yellow arrows). The T2* shortening is reflected in the lower tail of the corresponding histogram, where the uncorrected and the corrected results overlap.

**Figure 3:**
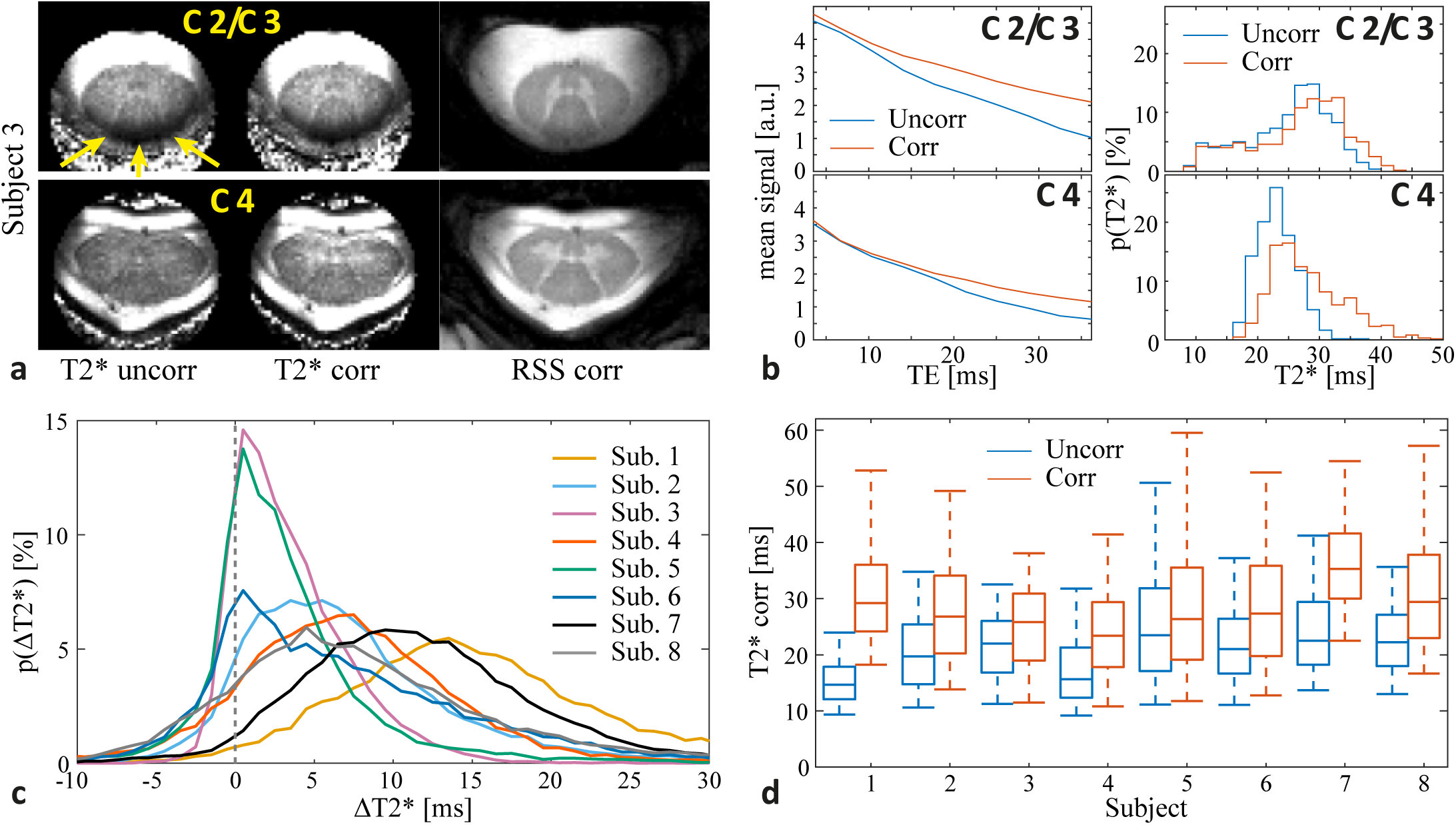
T2* quantification results from one subject (top) and a summary over all subjects (bottom). **a)** T2* maps without and with correction, next to the corrected RSS image, in two different slices from one subject. The arrows point to a band of low T2* due to static field inhomogeneity. **b)** Mean signal decay (left), and a histogram of the measured T2* values (right) inside the spinal cord mask without and with correction, for the two slices shown in a. **c)** Histogram of the voxel-wise difference in the measured T2* without and with correction 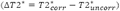 for all subjects. **d)** Box plot of the T2* distribution without and with correction inside the spinal cord mask for all subjects. The center line represents the median value, the box shows the 25^th^ and 75^th^ percentiles, and the whiskers show the 5^th^ and 95^th^ percentiles.

Figure 4 shows the multi-shot EPI time series mean image, standard deviation and tSNR in an axial slice at mid-vertebral level C5 in three subjects, without and with correction. The field fluctuations primarily cause data inconsistency between the different shots, leading to ghosting. The correction visibly reduced ghosting in all subjects. The magnitude of the ghosting varied considerably between subjects, as well as between volumes in the time series of a single subject. The time-varying ghosting leads to higher standard deviation and reduced tSNR over the time series. The correction improved the tSNR inside the spinal cord in all subjects. Figure 5 shows a sagittal view of the tSNR without and with correction in three subjects (for completeness, we show the subjects not included in Figure 4). The improvement was larger towards lower levels of the spinal cord, where the field fluctuations are stronger. The mean tSNR within the whole spinal cord mask increased by 32% on average over the subjects (range 6-59%), and the tSNR at around level C6 increased by on average 69% (range 10-135%) (Fig. 5b).

**Figure 4:**
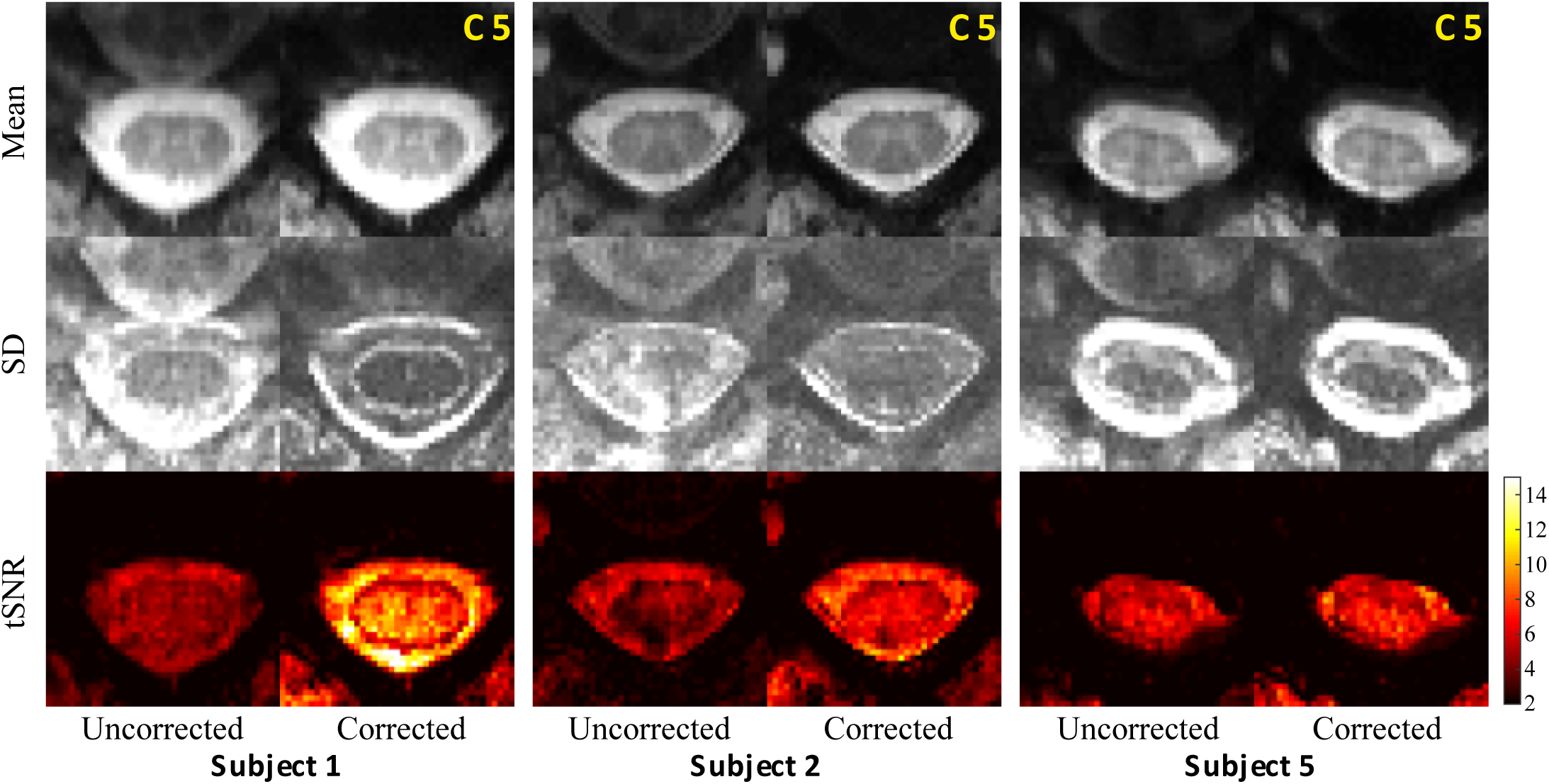
Multi-shot EPI mean image, standard deviation (SD) and tSNR in a single axial slice at C5 mid-vertebral level, without and with correction in three different subjects.

**Figure 5:**
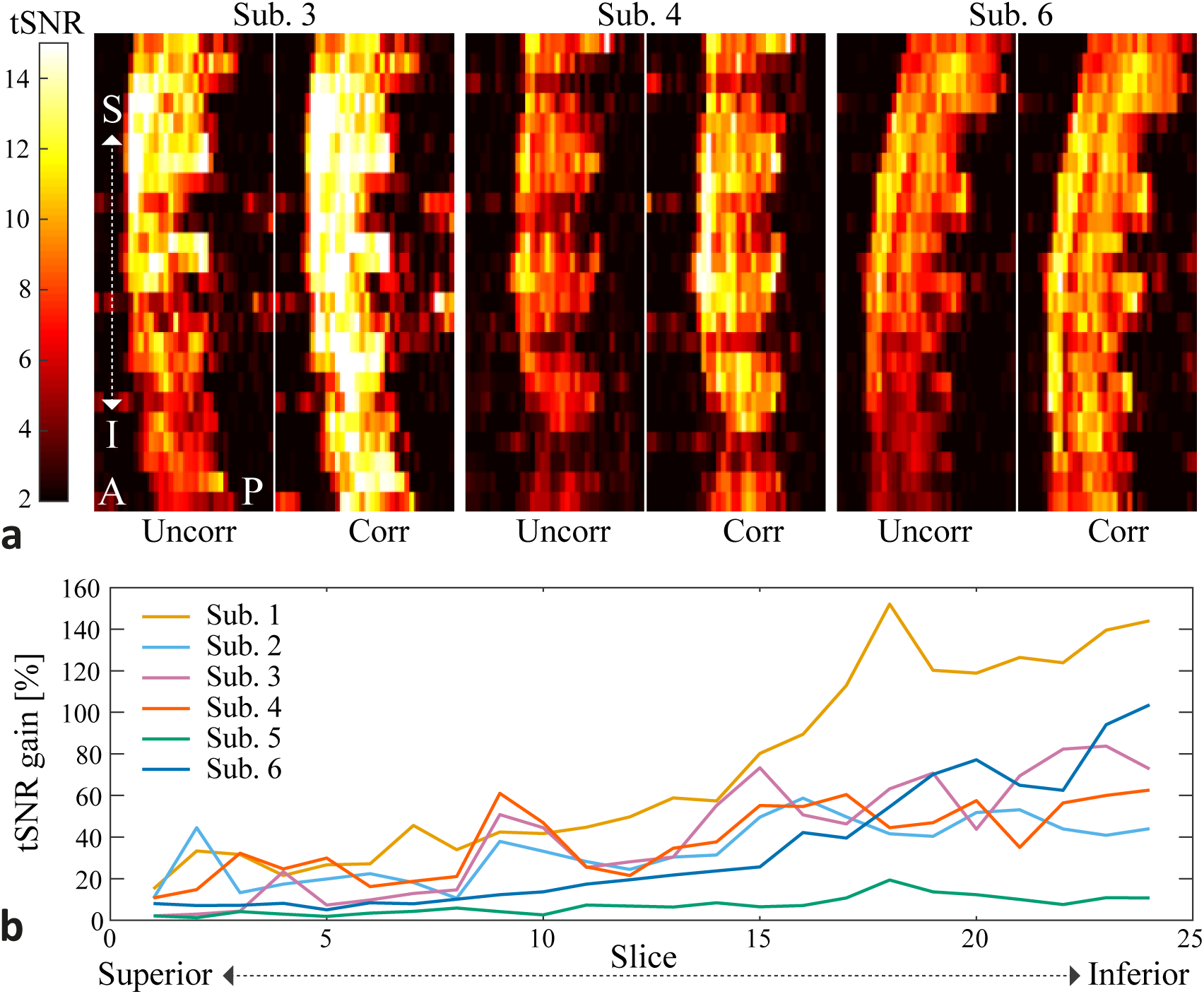
**a)** Sagittal tSNR maps for the multi-shot EPI without and with correction in three different subjects. **b)** Relative tSNR gain due to the correction for all subjects. The tSNR was averaged over all voxels inside the spinal cord mask in each axial slice (slice numbers counted superior to inferior, covering approximately C2-C6).

## Discussion

In this work, we have presented a correction for breathing-induced field fluctuations in spinal cord T2*-weighted imaging. The correction utilizes a respiratory trace to remove breathing-related phase instabilities in the acquired MR signal, and can reduce ghosting in multi-shot anatomical and functional acquisitions. The method relies on a short calibration scan, but does not require any additional hardware beyond a respiratory bellows for tracking the breathing state. Dynamic B0 fields are a considerable challenge in ultra-high field spinal cord imaging (17), and the method could therefore help to make greater use of the benefits of higher field strengths.

Quantitative T2* mapping was one of the demonstrated use cases for the proposed correction. In the uncorrected case, the T2* values in the spinal cord were systematically underestimated. This happens because the phase offset corresponding to a given field offset scales with the echo time, leading to increased ghosting in later echoes. The ghosting scatters the signal from the spinal cord over the image, thereby mimicking T2* signal loss inside the cord. The measured T2* values with correction were in good agreement with values previously reported at 7T (18). The previous study used a navigator echo for phase stabilisation (18,19). Navigators, however, take up time in the sequence and may prolong the minimum achievable TE and/or TR. In the context of T2* mapping, a navigator readout either replaces the first imaging echo or is appended after the last. In the former case, the early part of the decay, which carries information about the proton density of the tissue, is lost. In the latter case, at long TE the navigator may have too low SNR for robust phase extraction, which may produce an unstable correction that can even increase artefacts. Both cases limit the feasible echo times of the imaging readouts, and our navigator-less protocol is thus able to extend the observable part of the T2*-decay. Measuring the full decay provides more information for the exponential fit and could potentially allow for more complicated signal models as compared to a mono-exponential decay.

The second application investigated in this work was multi-shot EPI time series for functional imaging. In brain imaging, functional acquisitions are routinely performed with single-shot EPI. Single-shot EPI is relatively insensitive to time-varying field offsets, which translate into apparent motion in the time series that can be addressed retrospectively with motion correction algorithms. However, fMRI of the spinal cord at 7T has to date been conducted with 3D multi-shot EPI acquisitions (20,21), to reduce distortion and signal dropout due to local static field inhomogeneity. Multi-shot sequences are more susceptible to time-varying field effects, as this causes phase inconsistencies between shots. We here demonstrated that the tSNR of multi-shot EPI can be improved with correction for the breathing-induced fields, especially towards lower levels of the cervical spinal cord. This may improve the sensitivity to detect neural activation in fMRI of the spinal cord, especially in combination with post-processing methods to further reduce the impact of signal variations of physiological origin (22).

The proposed correction method was able to greatly reduce ghosting artefacts, but did not completely eliminate them. Residual artefacts after correction are also frequently observed with phase navigators. A number of potential reasons for incomplete correction can be identified. Firstly, the proposed correction relies on a reproducible and linear relationship between respiratory trace signal and the field state. This is a good approximation during regular shallow breathing, but is less reliable during deep or irregular breathing (5). Secondly, the spatial field profile of the time-varying fields may not be perfectly homogenous within the transversal slice. Previous characterizations of breathing-induced fields in the cervical spine have demonstrated a field gradient in the anterior-posterior direction in slices through the center of the neck, and a highly non-linear field component in slices closer to the thorax (5). Thirdly, the correction only accounts for breathing-induced fields, and not for actual motion of the subject nor for field variations from other sources. Slight actual motion of the neck associated with the breathing cycle is expected. Furthermore, swallowing induces both local motion of tissue and field variations of up to about 40 Hz (5). Residual ghosting due to swallowing was occasionally observed in the multi-shot EPI time series.

The trace-based correction method could potentially be expanded to account for the above additional perturbations. For example, slice-dependent anterior-posterior field gradients could be estimated from the sagittal FLASH images, and the phase encoding gradient could be adjusted accordingly in the reconstruction. A non-linear model linking the trace to the field variations may be able to capture a larger range of fluctuations and breathing modes. Furthermore, additional external devices,such as NMR field probes (23) or motion-tracking optical devices (24) could yield more information about both the field state and the motion, which could then be incorporated into the reconstruction model (25). A further improvement to the current implementation of the correction method would be to eliminate the manual steps in the processing of the field calibration data, i.e. the spinal cord mask creation and the exclusion of time points affected by swallowing. This would be a crucial step to allow for integration into standard acquisition protocols. Potentially this could be achieved with the Spinal Cord Toolbox (16) for masking, combined with automatic outlier detection in the field time-courses.

## Acknowledgments

The authors would like to thank Nadine Graedel for software contributions for the static phase correction. This project has received funding from the Wellcome Trust (Strategic Award 102645/Z/13/Z, Senior Research Fellowship 202788/Z/16/Z) and from the European Union’s Horizon 2020 research and innovation programme under the Marie Sklodowska-Curie grant agreement No 659263. The Wellcome Centre for Integrative Neuroimaging is supported by core funding from the Wellcome Trust (203139/Z/16/Z).

